# Functional anomaly mapping reveals local and distant dysfunction caused by brain lesions

**DOI:** 10.1101/464248

**Authors:** Andrew T. DeMarco, Peter E. Turkeltaub

## Abstract

The lesion method has been a cornerstone in the endeavor to understand brain-behavior relationships in humans, but has relied on the flawed assumption that anatomically abnormal tissue functions abnormally and anatomically normal tissue functions normally. To address this longstanding problem, we introduce an approach to directly map the degree of functional anomaly throughout the brain in individual patients. These functional anomaly maps identify anatomical lesions and are stable across measurements. Moreover, the maps identify functionally anomalous regions in anatomically normal tissue, providing a direct measure of remote effects of lesions such as diaschisis. Lesion-behavior mapping using these maps replicates classic behavioral localization and identifies relationships between tissue function and behavior distant from the anatomical lesions. This method provides brain-wide maps of the functional effects of focal lesions, which could have wide implications for one of the most important methods in neuroscience.

## Introduction

Lesion-behavior relationships have been recognized since ancient times (Helgason, 1987). Identifying consistent relationships between lesion location and behavioral deficits were keys to the localizationist debates of the 19^th^ century (Bartolomeo, 2011; Dronkers et al., 2007; Steinberg, 2014). To this day, lesion studies remain a cornerstone of neuroscience because they permit a strong causal inference that recently popular methods like functional MRI do not (Adolphs, 2016; Fellows et al., 2005; Price et al., 1999; Rorden and Karnath, 2004). Contemporary lesion-behavior mapping incorporates modern neuroimaging methods to precisely map brain regions in which anatomical lesions cause behavioral deficits (Bates et al., 2003; Rorden et al., 2007).

Despite the importance of the lesion method, it has always suffered from a major drawback: it relies on the assumption that tissue that appears anatomically abnormal functions abnormally, and that tissue that appears anatomically normal functions normally. The first issue causes problems because lesions appear heterogeneous and often do not have clear boundaries on scans, so manual segmentation requires a number of arbitrary decisions. For the same reason, automated lesion segmentation methods perform inadequately in general (Maier et al., 2015). Most significantly, one cannot know whether tissue that appears somewhat abnormal anatomically is functioning normally or not, nor can one know the degree of dysfunction if present.

More importantly, anatomical lesion studies make the flawed assumption that tissue that appears anatomically normal is functioning normally. Yet, it has been recognized since the early 20^th^ century that lesions cause distant functional disturbances, a phenomenon known as diaschisis (Carrera and Tononi, 2014; Feeney and Baron, 1986; von Monakow, 1914). Modern ideas about brain function emphasize the role of large-scale networks (Bullmore and Sporns, 2009; Farah, 1994; Mesulam, 1990) and disruption of brain networks is thought to form the basis of diaschisis (Feeney and Baron, 1986; Grefkes and Fink, 2014; Price et al., 2001).

Positron emission tomography (PET) has detected metabolic disturbances distant from the lesion even in the chronic period (Metter, 1991), but this technique is not common because of its use of radioactive tracers. A number of studies have also used task-based fMRI to demonstrate functional disturbances distant from the lesion (Corbetta et al., 2005; Saur et al., 2006; Ward et al., 2003), but such task-based approaches are limited because they only measure dysfunctional neural processes associated with one specific experimental paradigm (Carter et al., 2012). More recent work has relied on resting state fMRI connectivity (Boes et al., 2015; He et al., 2007; Nair et al., 2015; Park et al., 2011; Yourganov et al., 2018), or hemodynamic lag at rest (Zhao, Lambon Ralph, & Halai, 2018) to examine remote effects of lesions, but these approaches have not yielded maps of brain-wide functional integrity in individual patients.

A method to directly map functional aberrations throughout the brain would obviate the arbitrariness of lesion segmentation by grading the degree of functional irregularity in anatomically abnormal regions. Further, this would extend lesion-behavior mapping beyond the boundaries of the anatomical lesions, providing a direct measure of functional differences between individuals related to diaschisis, compensatory plasticity, or individual differences that explain resilience to stroke.

We have developed a method that uses machine learning on 4D resting state fMRI data to map the degree of functional aberration throughout the brain in individual patients. We demonstrate that these functional anomaly maps (FAMs) identify the region of obvious anatomical damage, and that the maps are reproducible over time. We further demonstrate that the local degree of functional aberration in the unlesioned hemisphere relates to the degree of dysfunction in the roughly homotopic location of the lesioned hemisphere, strongly suggesting an effect of the lesion on the function of distant, seemingly intact, tissue. Finally, we demonstrate that lesion-behavior mapping using FAMs recapitulates the results of lesion-behavior mapping based on anatomical lesions, and importantly that it can identify regions outside the distribution of anatomical lesions in which functional aberration relates to behavioral outcomes. This method for direct brain-wide mapping of functional lesions could have wide implications for one of the most important approaches to neuroscientific inquiry.

## Results

### Functional Anomaly Maps Detect Anatomical Lesion Location

An anatomical lesion overlap map of the patients is shown in **Figure 1**. Of the 132 resting state datasets across 50 subjects for which FAMs were estimated, 128 (96.9%) were successful. Four failed to converge or contained zero support vectors with our hyperparameter values and control group. **Figure 2A** shows FAMs for an example patient over three scan sessions, and illustrates that FAMs contain a grading of functional anomaly at every voxel in the brain. **Figure 2B** shows FAMs for a sample of four patients. Although for the reasons outlined in the introduction the anatomical lesion masks cannot serve as a gold standard against which to test FAMs, we still predicted that in general, FAMs should identify the region of gross anatomical abnormality. Upon visual inspection, the anatomical lesion tracings (**Figure 2A** left column, red outline) appear very similar to areas of high functional anomaly load for each of three scan sessions (**Figure 2B**, right). In some cases, FAMs show areas of highest functional anomaly load outside the anatomical lesion. An example of this is shown in **Figure 2B**, in which a patient with a lesion in the deep white matter and basal ganglia of the left hemisphere (left pane, arrow) exhibits widespread abnormalities in the cortex of the left hemisphere. This pattern of mild metabolic changes throughout ipsilesional hemisphere has been previously observed on PET after some subcortical lesions (Metter, 1991). Although the FAMs in Figure 2B and 2C are thresholded to visually highlight areas of highest signal, at no point in our analyses is a threshold applied to FAMs. FAMs for all patients at all available time points are shown in **Supplemental Figure 1**.

**Figure 1.**
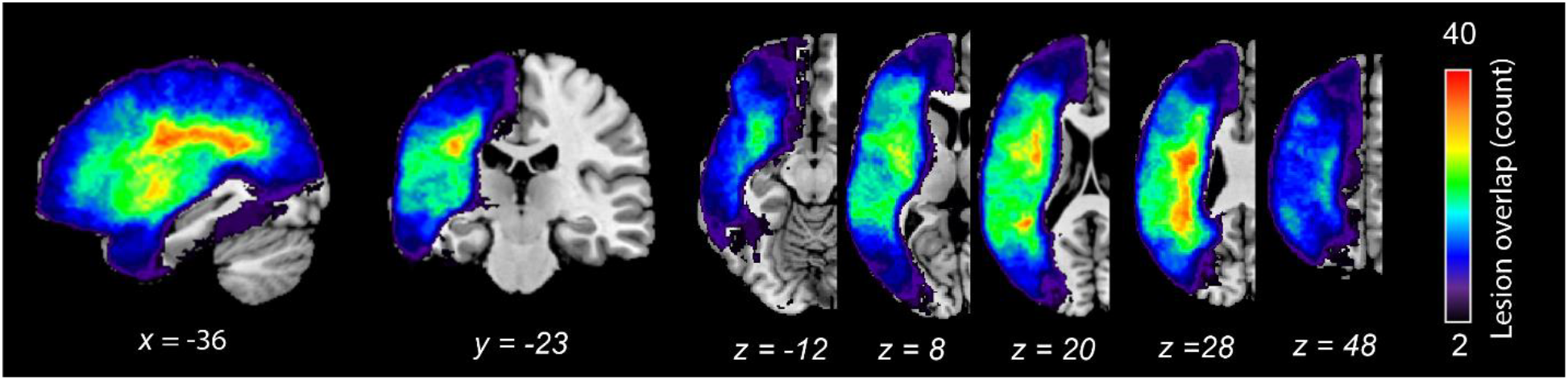
Overlap of anatomical lesions from the patient cohort.

**Figure 2.**
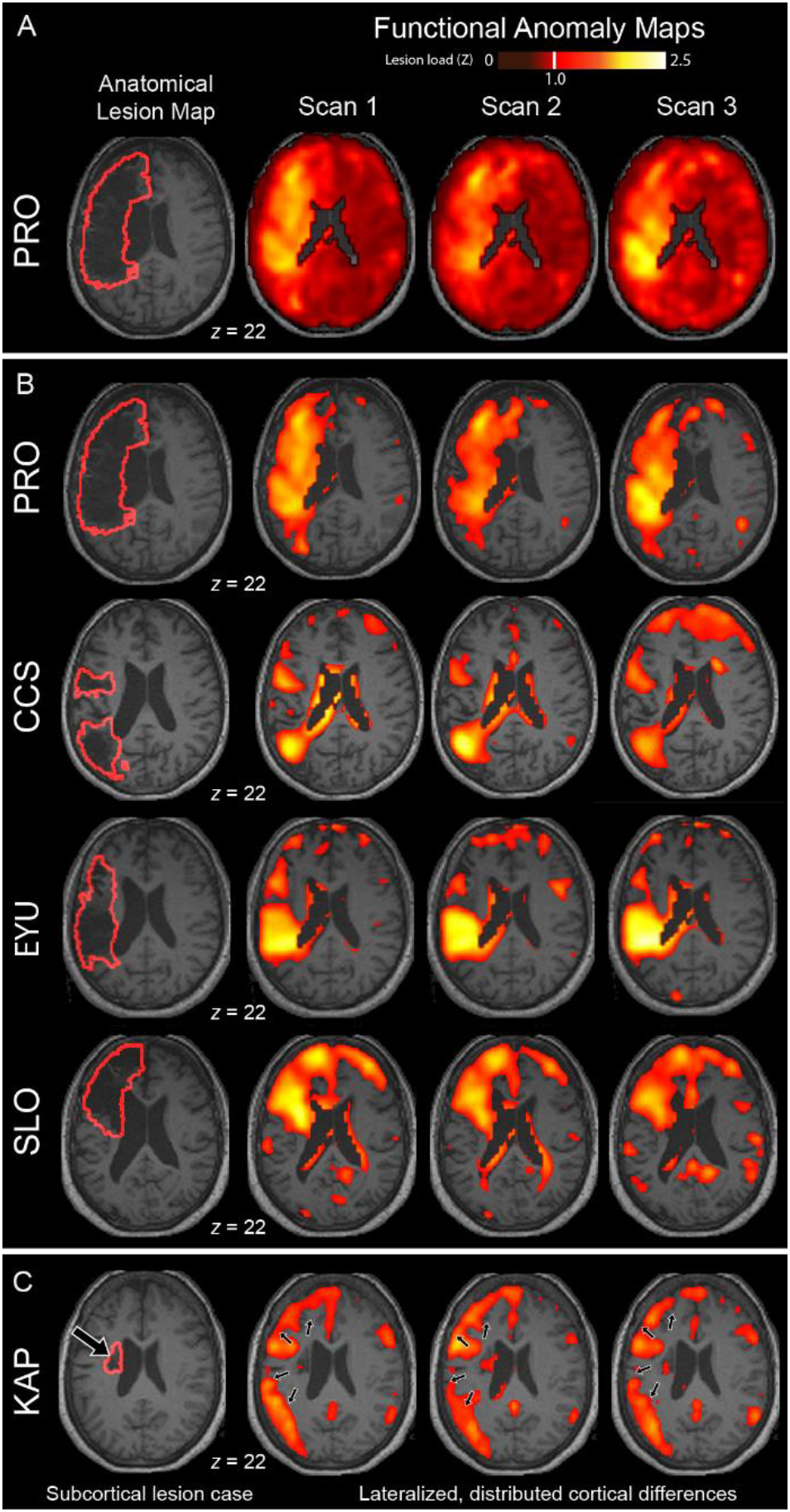
Examples of patients (rows) with anatomical lesion tracing outlined on their high-resolution anatomical scans (leftmost column, red outline) and FAMs (hot scale) for three scan sessions (columns). (A) Example FAM showing that all voxels in the brain are graded for statistical deviation relative to the control group. (B) An illustrative threshold is applied to FAMs for four example patients to highlight that in many individuals, the regions of greatest functional anomaly roughly correspond to obvious anatomical lesions. (C) An illustrative threshold is applied to another example patient’s FAMs to highlight that some patients with subcortical lesions (left, larger arrow) reliably showed areas of greatest signal throughout the left-hemisphere cortex in the lesioned hemisphere (small arrows). FAMs in panes (B) and (C) are thresholded at a functional anomaly load cutoff of *Z* > 1 to illustrate areas of highest signal.

We performed three analyses to determine the degree to which FAMs are sensitive to the left-hemisphere anatomical lesions in our patient cohort. First, we predicted that larger anatomical lesions would be associated with greater left-hemisphere functional anomaly load. As predicted, in a bivariate correlation, larger anatomical lesions were associated with greater average left-hemisphere signal in the FAMs (*r* = .49, *P* < .001) (**Figure 3A**).

**Figure 3.**
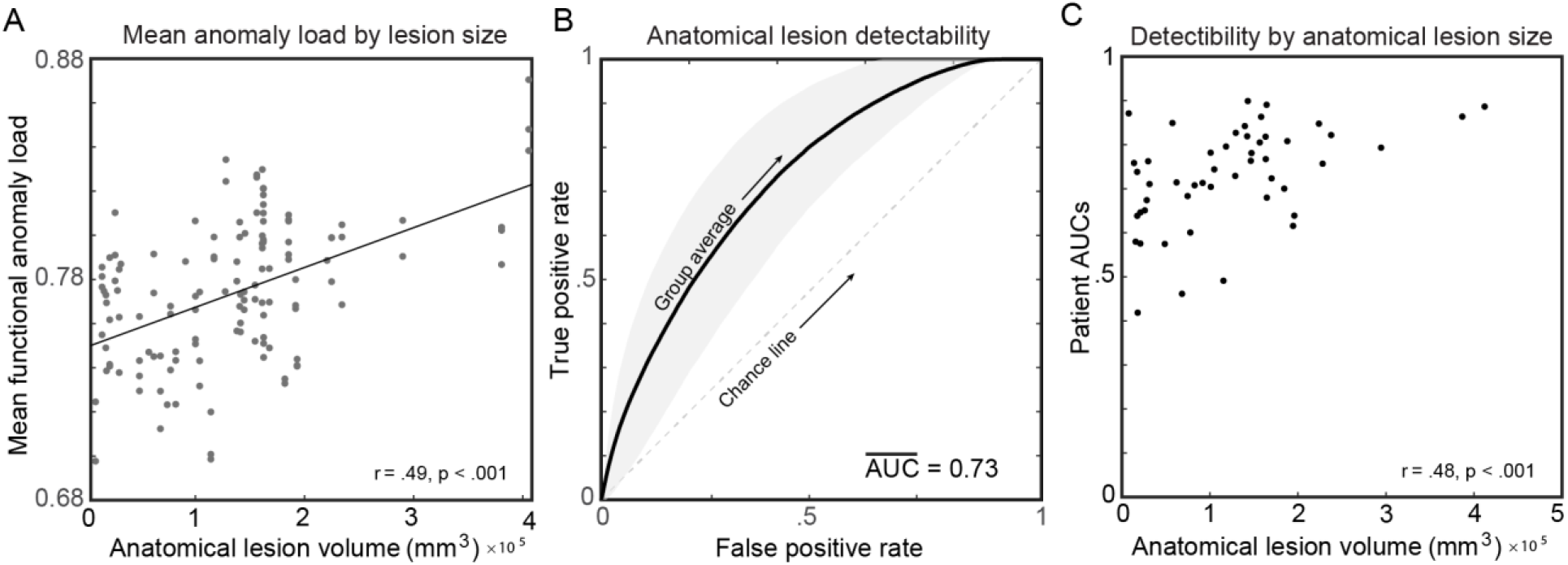
FAMs detected manually-traced anatomical lesions as evidenced by (A) a significant positive relationship between anatomical lesion volume and average function anomaly load within the left-hemisphere, (B) a group average ROC curve for the ability of FAMs to classify tissue labeled anatomically lesioned from background, and (C) a significant positive relationship between anatomical lesion volume and FAM classification accuracy for anatomical lesion maps.

Second, we predicted that the signal in FAMs would discriminate regions of anatomical lesion from unlesioned tissue. We tested this using a receiver operating characteristic (ROC) curve analysis in which we calculated area under the curve (AUC) for a patient’s FAM to accurately classify voxels of their anatomical lesion map. Average AUC was .73 (*SD* = .11), significantly better than a chance-level AUC of .5 (*t*(48) = 14.34, *P* < .001), indicating that the FAMs distinguished regions of anatomical lesion from unlesioned tissue (**Figure 3B**). AUC correlated with anatomical lesion volume (*r* = .477, *P* < .001), such that the larger the anatomical lesion, the more closely the location of high FAM signal corresponded to the location of the anatomical lesion (**Figure 3C**). ROC curves and AUCs for each patient are shown in **Supplemental Figure 2**.

Third, since one important potential use of FAMs is in lesion-behavior mapping analyses, we tested if lesion-behavior maps based on FAMs could localize scores with a ground-truth anatomical localization. Scores were derived for each patient as anatomical lesion load in each of 132 left-hemisphere atlas parcels. The FAM-based lesion-behavior maps localized the scores generated from atlas parcels to the source anatomical lesion location with a median distance of 2.01 cm (*SD* = 1.00 cm), significantly better than chance based on permutation analysis (*Z* = 10.6, *P* < .001; **Figure 4A**). The precision of localization using this approach is limited by lesion covariance and the size of the atlas parcels, so for comparison, we ran the same analyses using the anatomical lesions to localize scores based on the same anatomical lesion data. These lesion-behavior maps localized the scores generated from atlas parcels with a median distance of 1.95 cm (*SD* = 0.96 cm; **Figure 4B**). Localization precision using FAM data was not significantly different from the results using anatomical lesion data (*Z* = 0.71, *P* = .48).

**Figure 4.**
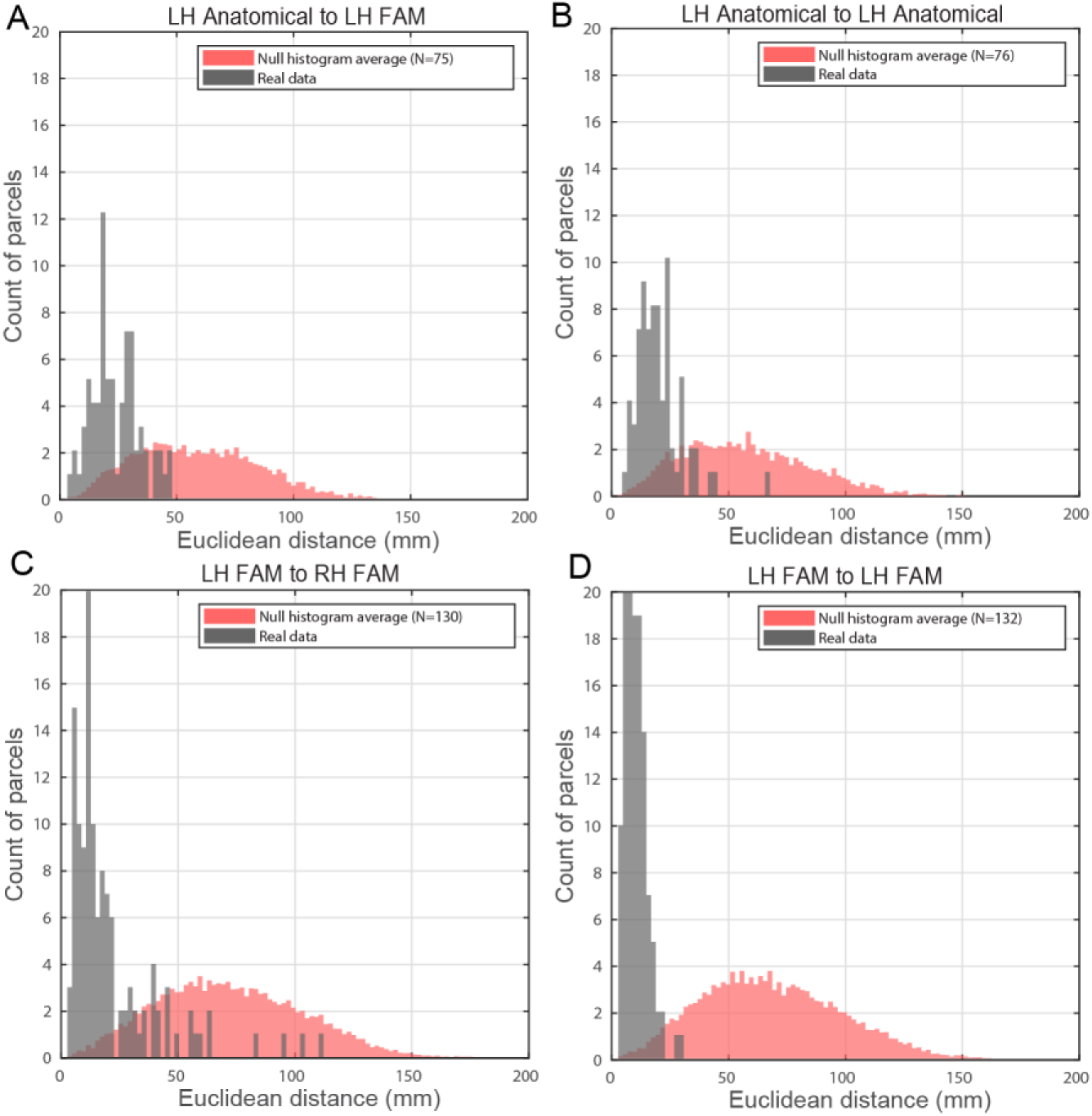
Accuracy of using SVR-LSM of FAMs to localize sources of scores that have ground-truth localization. The top row shows localization accuracy of scores with ground-truth localization derived from anatomical lesion load within left-hemisphere atlas parcels: (A) localization accuracy using left-hemisphere FAMs to map scores and, for comparison, (B) localization accuracy using left-hemisphere anatomical lesions to map scores. The bottom row shows localization accuracy of scores with ground-truth localization derived from FAM values in left-hemisphere atlas parcels: (C) homotopic localization accuracy of mapping using right-hemisphere FAMs and, for comparison, (D) localization accuracy of mapping using left-hemisphere FAMs. For each analysis, a histogram of localization accuracy is shown for the real data (gray) and for comparison a histogram of the null distribution (red). All histograms have a bin width of 2. Distances are computed from center-of-mass between atlas seed parcel used to generate the score to that of the resulting SVR-β map thresholded at >7.

### Functional Anomaly Map Signal Indexes Functional Integrity of Spared Tissue

Next, we tested if transcallosal diaschisis (Feeney and Baron, 1986; von Monakow, 1914) would be measurable as functional aberrations in the unlesioned hemisphere. To test this question, we generated scores based on the mean FAM signal in atlas parcels placed in the lesioned left-hemisphere and performed lesion-behavior analyses to localize right-hemisphere regions in which the FAM signal related to these left-hemisphere scores. We reasoned that the FAM signal in the unlesioned right-hemisphere should relate to the degree of functional anomaly in the homotopic areas of the lesioned left-hemisphere. Confirming our prediction, the right-hemisphere regions identified by these analyses had a median distance of 1.28 cm (*SD* = 1.97 cm) from the homotopic site of the corresponding left-hemisphere regions (**Figure 4C**), significantly better than chance based on permutation testing (*Z* = 12.9, *P* < .001). To test the upper bound of precision for this analysis, we then used the same approach to localize regions in left-hemisphere FAMs in which the signal related to the scores derived from the same left-hemisphere FAMs. The median localization precision was 0.82 cm (*SD* = 0.47 cm; **Figure 4D**).

### Functional Anomaly Maps Are Reproducible Over Multiple Independent Sessions

For the 36 patients for whom data from multiple sessions was available, the average intraclass correlation coefficient (ICC) for FAM values across the brain ranged from .66 to .89 (*M* = .80, *SD* = .06; **Figure 5A**), in the good-to-excellent range based on a commonly cited qualitative interpretation of ICC (Bennett and Miller, 2010; Cicchetti and Sparrow, 1981). For each of these patients, average voxelwise correlation between pairs of runs ranged from .68 to .89 (*M* = .80, *SD* = .06; **Figure 5B**, **Figure 5C**).

**Figure 5.**
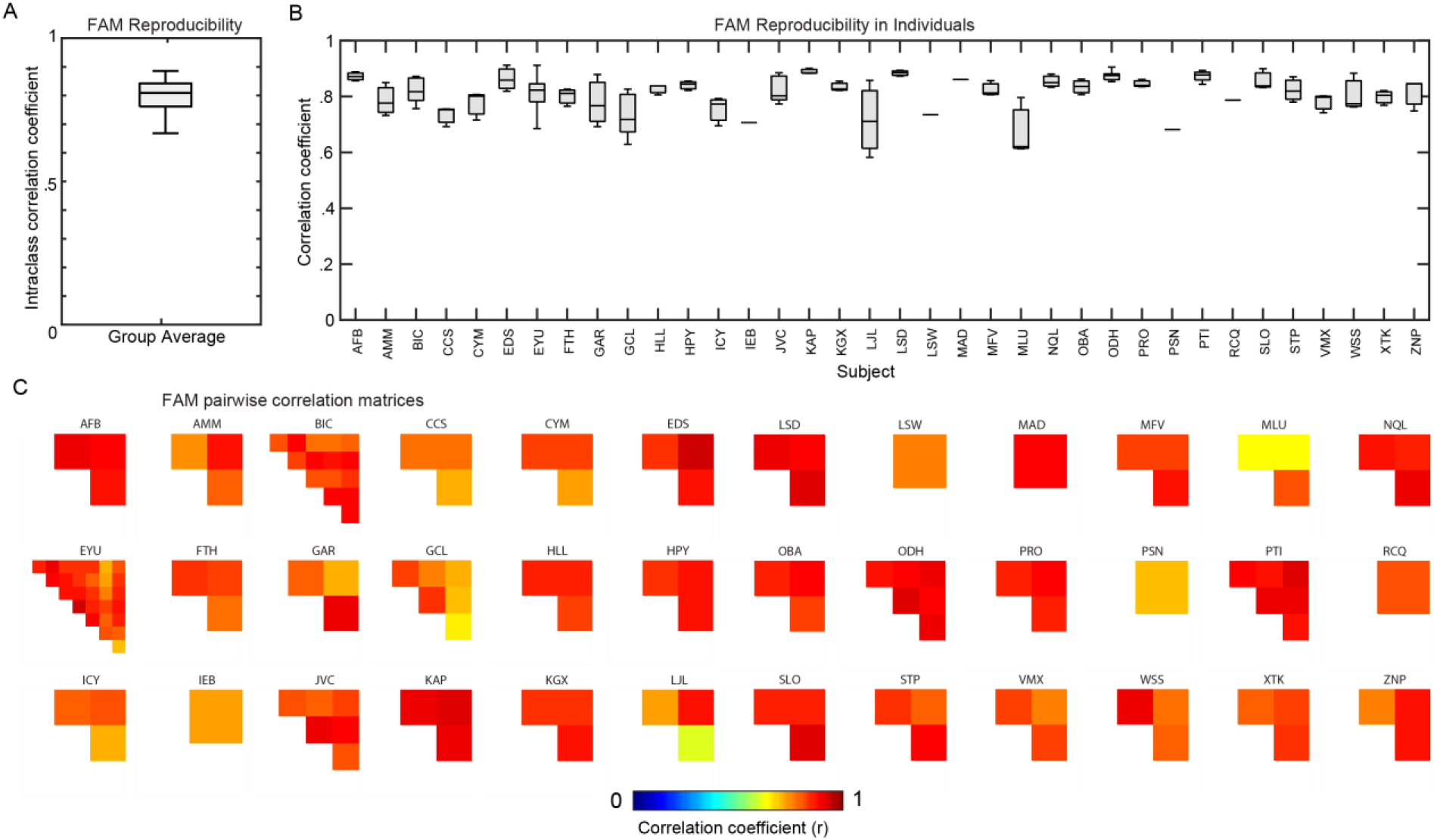
Reproducibility of FAMs for the 36 subjects with data available from multiple scanning sessions. (A) A boxplot of whole-brain voxelwise intraclass correlation coefficient across the group. (B) Average whole-brain voxelwise correlation coefficients between pairs of resting state runs for individual patients. (C) Voxelwise correlations (jet scale) between pairs of resting state runs for individual patients.

### Functional Anomaly Map Signal Relates to Behavior In Both Lesioned and Unlesioned Tissue

One of the main potential uses for FAMs is for lesion-behavior mapping analyses. To assess their utility for this purpose, we performed multivariate lesion-behavior mapping analysis using the FAMs (called functional lesion-behavior mapping below) and comparable analyses using the anatomical lesion maps (called anatomical lesion-behavior mapping below), examining four behaviors commonly affected by left-hemisphere stroke: auditory comprehension, phonological processing, speech fluency, and right-hand strength.

Shown in **Figure 6A**, functional lesion-behavior mapping revealed that impaired auditory comprehension was associated with a single cluster encompassing anterior/middle left temporal lobe, sagittal stratum, Heschl’s gyrus, left posterior insula and external capsule (*P* < .001, 48768 mm^3^). Anatomical lesion-behavior mapping produced a single cluster encompassing superior temporal gyrus (*P* = .002, 31812 mm^3^), largely overlapping the functional results. The main differences were that functional results included the left lateral/medial middle and inferior temporal gyri and left anterior fusiform gyrus, while anatomical results were largely restricted to the superior temporal gyrus, but extended into the posterior superior temporal gyrus. These findings are consistent with the classic localization of auditory comprehension to the temporal lobe, as shown by prior stroke lesion studies (Bonilha et al., 2017; Hillis et al., 2017) and fMRI studies of healthy controls (Price, 2012). The fact that the functional results extend more ventrally into the temporal lobe likely relates to poor anatomical lesion coverage in that region.

**Figure 6.**
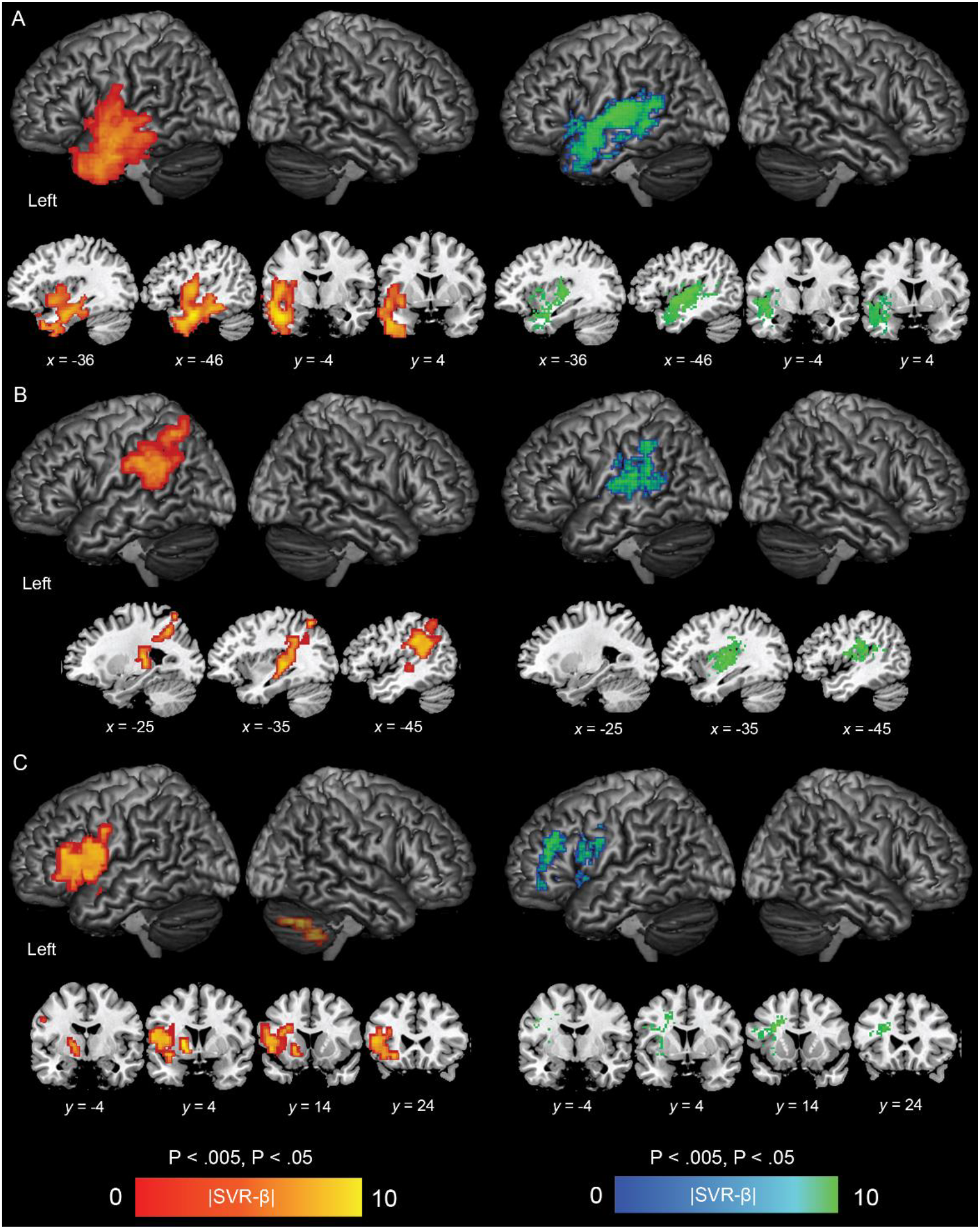
Lesion-behavior mapping results for three speech and language behaviors including (A) auditory comprehension, (B) phonology, and (C) speech production. Results are shown for FAM data (left, red-yellow scale) and traditional anatomical lesion data (right, blue-green scale). With the exception of the right cerebellar findings for functional lesion-behavior mapping for speech production deficits, which didn’t quite reach full correction for multiple comparisons (translucent), all results are shown as the absolute value of SVR-β values thresholded voxelwise at *P* < .005 and corrected for multiple comparison by cluster extent at *P* < .05 based on 10,000.

Shown in **Figure 6B**, functional lesion-behavior mapping revealed that impaired phonological processing was associated with a single cluster encompassing the left supramarginal gyrus, neighboring intraparietal sulcus, and underlying white matter (*P* = .001, 19648 mm^3^). Anatomical lesion-behavior mapping revealed a single cluster (*P* = .007, 16281 mm^3^), which appeared patchy around the left temporoparietal junction near the strongest functional results, and extended medially and anteriorly into posterior insula, parietal operculum, and posterior external capsule. Both analyses converged at the temporoparietal junction. This localization is consistent with modern neuroanatomical language models that implicate this region as the locus for converting auditory input to phonological representations (Hickok and Poeppel, 2007) and patient studies which find that damage to this area relates to repetition deficits that characterize conduction aphasia (Buchsbaum et al., 2011; Hickok and Poeppel, 2000; Hickok et al., 2003).

Shown in **Figure 6C**, functional lesion-behavior mapping found that impaired speech fluency was associated with a cluster encompassing ventral premotor cortex, pars opercularis and triangularis, anterior corona radiata, anterior insula and neighboring external capsule, and the anterior limb of the internal capsule (*P* = .001, 21120 mm^3^). A second cluster in right cerebellum approached significance (*P* = .098). Anatomical lesion-behavior mapping found impaired speech fluency was associated with a single patchy cluster of anatomical damage including ventral premotor cortex, pars opercularis, external capsule, a larger swath of anterior corona radiata and some superior corona radiata, extending into inferior frontal sulcus and the middle temporal gyrus (*P* = .009, 13312 mm^3^). The main difference between the two analyses was that the functional results showed larger cortical findings and included anterior insula, while the anatomical results included more anterior corona radiata and extended into middle frontal sulcus/gyrus. The functional results in right cerebellum were naturally absent from the anatomical results. The left frontal lobe localization is consistent with prior stroke lesion studies (Basilakos et al., 2015; Graff-Radford et al., 2014; Itabashi et al., 2016) and work on patients with progressive apraxia of speech (Josephs et al., 2006). In addition, speech motor planning is among the functions to which the right cerebellum is thought to contribute (Mariën et al., 2014; Stoodley et al., 2012).

Finally, shown in **Figure 7**, functional lesion-behavior mapping revealed three left-hemisphere clusters associated with right-hand strength that failed to reach significance based on cluster-level correction for multiple comparison. Two of these clusters overlapped with primary motor cortex including the hand area (*P* = .07, 6016 mm^3^), descending corticospinal tract including superior corona radiata, and the posterior limb and retrolenticular portion of the internal capsule (*P* = .12, 4480 mm^3^). Together these two clusters overlap the canonical motor system, but were split into two at the fibers of the descending corticospinal tract. The third cluster involved frontal white matter (*P* = .063, 6400 mm^3^) including the anterior superior corona radiata and underlying body of the corpus callosum. One right-hemisphere cluster in the ventral premotor cortex, not including underlying white matter, survived correction for multiple comparisons (*P* = .02, 10304 mm^3^). Anatomical lesion-behavior mapping produced a single significant left-hemisphere cluster encompassing primary motor cortex and descending corticospinal tract, overlapping with the functional results (*P* = .04, 5390 mm^3^). The main difference between the analyses was that anatomical results did not reach as far dorsally into cortex, but its higher spatial resolution allowed it to detect the slender descending corticospinal tract fibers without any breaks. In addition, the anatomical analysis naturally produced no right-hemisphere results, as anatomical lesion coverage was limited to the left-hemisphere in our cohort. These left-hemisphere findings are consistent with the canonical motor system architecture, with both methods converging on the descending corticospinal tract (MNI *x* = -24, *y* = -17) and overlapping the posterior limb of the internal capsule. The right-hemisphere region identified by functional analysis is convergent with evidence that contralesional premotor cortex plays a role in recovery after motor stroke, with meta-analytic neuroimaging results showing consistent recruitment of this area to support residual function in post-stroke motor deficits (Rehme et al., 2012). It is also consistent with findings that inhibitory neurostimulation over contralesional dorsal premotor cortex impairs recovered motor performance in patients with subcortical stroke (Lotze et al., 2006).

**Figure 7.**
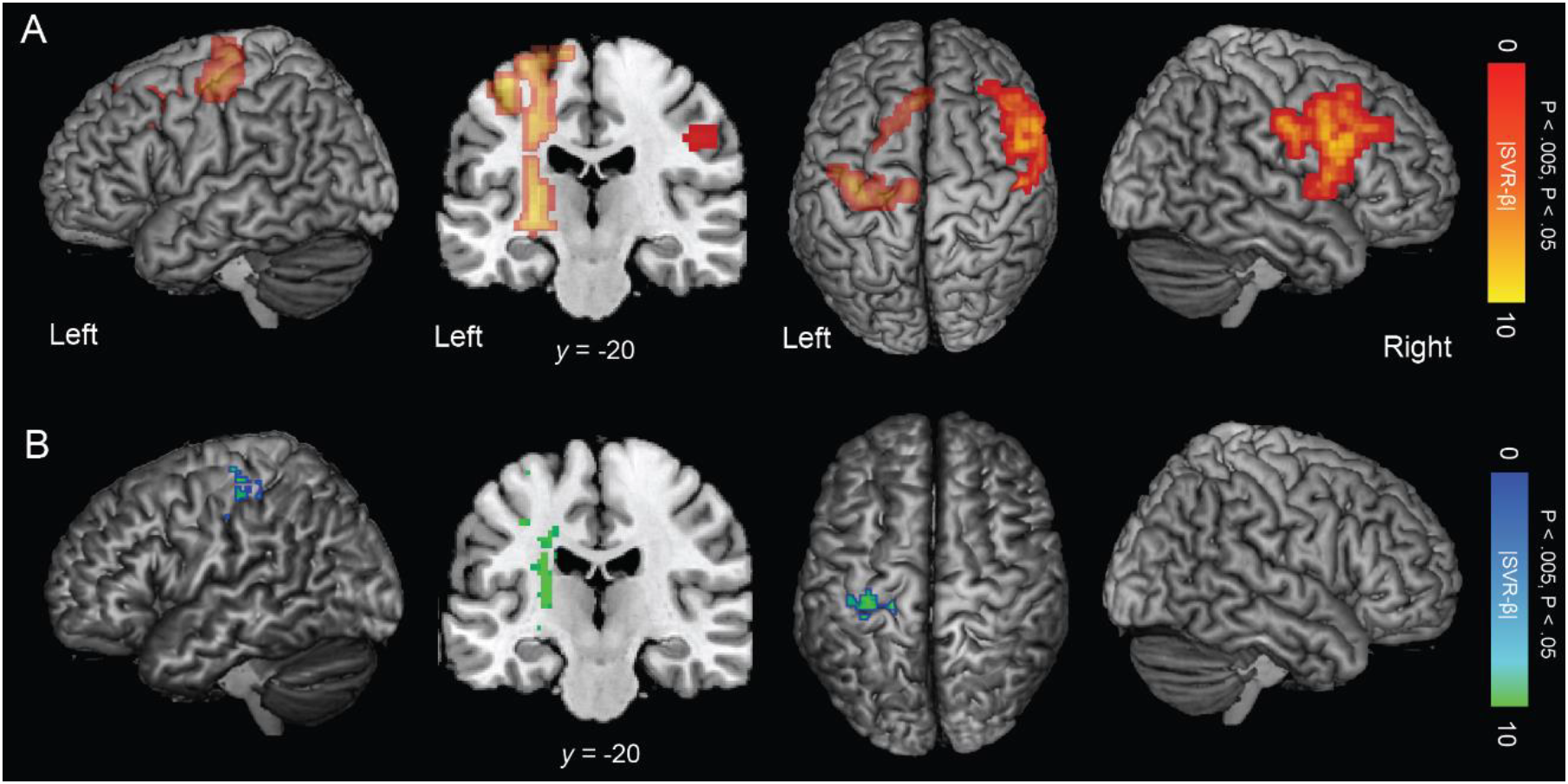
Results of lesion-behavior mapping for right-hand pinch strength using (A) FAMs (red-yellow scale) and (B) traditional anatomical lesion data (blue-green scale). With the exception of left-hemisphere functional lesion-mapping results which failed to quite reach full significance after correction for multiple corrections (translucent), results are shown as the absolute value of SVR-β values thresholded voxelwise at *P* < .005 and corrected for multiple comparison by cluster extent at *P* < .05 based on 10,000 permutations.

## Discussion

Lesion-behavior association methods have been limited by assumptions regarding the relationship between anatomical appearance of tissue and its functional integrity. Here, we have introduced a method to derive brain-wide maps of functional integrity from 4D resting fMRI data. These maps are replicable when derived from data acquired on different days, and are sensitive to the location of the anatomical lesion. Further, signal in the unlesioned hemisphere relates to the functional integrity of the homotopic locations in the lesioned hemisphere, suggesting that this signal is meaningful and may reflect transcallosal diaschisis. When used to localize behavioral deficits, these maps replicate classical patterns of behavioral localization, and are able to detect relationships between functional integrity and behavior in regions distant from the anatomical lesions. Together these findings demonstrate that this technique can reliably map behaviorally relevant functional integrity of tissue throughout the brain in individual stroke survivors. This new approach to lesion measurement could have widespread implications for one of the key methods of neuroscientific inquiry.

The long-standing approach to lesion identification based on anatomical appearance has come with critical limitations on attempts to interpret the consequences of lesions. In addition to the unclear relationship between anatomical signal and functional integrity mentioned above, defining lesion boundaries by anatomy alone requires arbitrary cutoffs, for example on the degree of white matter signal abnormality included in the lesion boundaries. Ultimately, anatomical lesion boundaries are unreliable because the tissue surrounding a lesion often exhibits metabolic or perfusion abnormalities not visible on anatomical scans (Forkel and Catani, 2018; Hillis et al., 2001; Karnath et al., 2018; Kuhl et al., 1980; Metter, 1991; Metter et al., 1981; Richardson et al., 2011). Identification of lesions based on anatomy alone also fails to detect distant dysfunction due to diaschisis (Carrera and Tononi, 2014; Feeney and Baron, 1986; Finger et al., 2004). Indeed, it is insensitive to any signal in residual tissue, including functional attributes of spared regions which may explain resilience to lesion effects (Abdullaev and Posner, 2005; He et al., 2007) and compensatory plasticity in spared regions which may contribute to behavioral outcomes (Corbetta et al., 2005; Nelles et al., 1999; Siegel et al., 2018). A method such as the one we present here addresses all of these issues. Although there is much room for further development of this technique, we clearly demonstrate that behaviorally relevant signal reflecting the functional status of tissue throughout the brain can be derived from a 4D BOLD time-series.

Previous approaches to identify functional abnormalities distant from lesions have also used rsfMRI, which has provided an avenue to explore intrinsic brain activity in the absence of a task (Snyder and Raichle, 2012). One approach, lesion network mapping (Boes et al., 2015), combines anatomical lesion information with functional connectivity, a common analytic approach to fMRI data that involves measuring long-range correlations in the spontaneous BOLD signal (Biswal et al., 1995). Functional connectivity has revealed a number of large-scale brain networks that correspond to cognitive states reliably across individuals (Fox and Raichle, 2007). Lesion network mapping takes advantage of these normative findings to infer the possible contribution of spared tissue to lesion-mapping results (Boes et al., 2015). By identifying a network of regions in a normative connectome that exhibit strong functional connectivity with each patient’s anatomical lesion, the method can deduce where areas of projected connectivity disruption might relate to behavioral symptoms. The approach has identified regions that explain an impressive diversity of specific symptoms associated with variable anatomical lesions sites (Darby et al., 2016, 2018; Fasano et al., 2017; Ferguson et al., 2017; Fischer et al., 2016; Laganiere et al., 2016; Sutterer et al., 2016). Still, this approach does not measure brain function directly in patients and relies on anatomically defined lesions. Connectome-based lesion-symptom mapping is another approach for understanding the contribution of spared tissue using structural or functional connectivity measured between brain regions. It has been proposed as a supplement to traditional anatomical lesion-symptom mapping’s poor sensitivity to white matter damage (Del Gaizo et al., 2017; Gleichgerrcht et al., 2017; Yourganov et al., 2016). When applied to structural data, this method finds the contribution of white matter disconnections to behavioral outcomes using a whole-brain structural connectome based on white matter tractography (Hagmann et al., 2008). Connectome-based lesion-symptom mapping has also been applied to functional connectivity matrices measured during resting state fMRI. This approach identifies locations where increased or decreased correlation between regions are associated with changes in behavioral deficits in post-stroke aphasia (Yourganov et al., 2018). Aside from functional connectivity, Zhao and colleagues (Zhao et al., 2018) associated regional hemodynamic lag or advance in spared tissue with specific behavioral deficits in post-stroke aphasia, but this method focuses on temporal hemodynamic disruption rather than the more general question of identifying areas of functionally aberrant tissue, and no attempt was made to validate this tool for mapping functional anomalies in individual patients.

The current method obviates the need for arbitrary anatomical segmentation of lesions, and is able to directly map the degree of voxelwise functional aberration throughout the brain in individual patients, identifying signals in both the area directly affected by the lesion and in anatomically spared regions. There are advantages to using FAMs specifically for lesion-behavior mapping. Lesion-behavior mapping using anatomical data has been challenged by modest sample sizes and variable lesion coverage, with both considerations limiting statistical power to detect lesion-behavior relationships (Inoue et al., 2014; Kimberg et al., 2007; Lorca-Puls et al., 2018). The use of FAMs allows participants to contribute multiple scans to a single lesion-behavior mapping analysis and ensures data in every voxel of the brain, meaning it could help with sensitivity for detecting lesion-behavior relations in regions where anatomical lesion coverage typically tends to be poor (e.g. ventral occipitotemporal gyrus). In addition, anatomical lesion-behavior mapping has been observed to have poor sensitivity for white matter damage (Gleichgerrcht et al., 2017), but the method presented here appears to show sensitivity to effects in both gray and white matter.

A key outstanding question regarding the FAM method is what biological processes the anomalous signal represents. The approach measures the degree to which the BOLD signal differs from control subjects over time. However, since aberrations in the ongoing BOLD fluctuations can produce either positive or negative differences from controls, the FAM signal has no sign. For this reason, the differences between a patient and control group identified by FAM could theoretically relate to either dysfunction or hyper**-**function, either occurring at a hemodynamic or a neural level. It is logical that the most extreme values would correspond to the area of obvious anatomical lesion, either due to gross disruption of hemodynamic response or due to gross neural dysfunction, and this is what we found. But signal elsewhere could reflect any difference in the patient relative to the control group. These differences could arise from negative biological effects of the stroke on surviving tissue, such as diaschisis (i.e., distant dysfunction due to network level disruption) (Carrera and Tononi, 2014; Feeney and Baron, 1986; von Monakow, 1914), hemodynamic differences (Bonakdarpour et al., 2007; Siegel et al., 2016; Zhao et al., 2018), maladaptive disinhibition (Allred et al., 2010; Ferbert et al., 1992; Johansen-Berg et al., 2002; Letzkus et al., 2015), or atrophy due to deafferentation of cell bodies in otherwise spared regions of the brain (Bonilha and Fridriksson, 2009; Bonilha et al., 2014). Other differences in anatomically intact brain regions could reflect negative behavioral consequences of stroke, such as learned disuse (Levin et al., 2009; Taub et al., 2006), or maladaptive behavioral compensation (Levin et al., 2009; Takeuchi and Izumi, 2012). Conversely, the signal could represent changes related to compensatory plasticity (Takeuchi and Izumi, 2013), including plasticity directly related to the biological effects of the stroke damage or plasticity related to environmental enrichment (e.g., therapy) (Biernaskie and Corbett, 2001; Döbrössy and Dunnett, 2001; Nithianantharajah and Hannan, 2006) or increased behavioral reliance on spared abilities (Bury and Jones, 2002; Cirstea and Levin, 2000; Luke et al., 2004; Rauschecker, 1995). Finally, the signal could relate to individual differences that predated the stroke (Hillis and Tippett, 2014; Nunnari et al., 2014; Pearson-Fuhrhop et al., 2012), although in a relatively homogeneous sample these would be expected to be smaller than the other effects.

These different potential sources of the FAM signal are difficult to disambiguate in individual patients at present, but clues regarding the prevailing sources of the signal can be gleaned from patterns across the group. For example, we found that the signal in the right hemisphere relates to the signal in homotopic areas of the left hemisphere, strongly suggesting a direct biological effect of the lesion, although the specific nature of that change remains unknown. We have demonstrated two examples of functional lesion-symptom mapping in which the contralesional functional integrity relates to behavioral outcomes. In these cases the inverse relationship between the signal and the behavior suggests that the anomalous signal measured there has negative effects on behavior and so likely reflects diaschisis, maladaptive disinhibition, or deafferentation atrophy rather than compensatory plasticity. Further research will be needed to investigate the nature of the signal, and to develop methods that can discriminate between the various potential causes of abnormal functional signals distant from the lesion.

Some limitations of relying on BOLD signal to detect lesions should be considered. Because this method relies on fMRI data, it is sensitive to head movement and has poorer spatial resolution than high-resolution anatomical scans. This is evident in our motor analysis results in which the descending corticospinal tract breaks into two clusters in the functional analysis because the fibers exceed the spatial resolution of our functional voxels. Moreover, although FAM provides whole-brain coverage for lesion detection, regional differences in sensitivity may occur due to hemodynamic differences between different tissue types or signal dropout and distortion from magnetic susceptibility differences at air tissue interfaces as with a typical fMRI analysis (Caparelli et al., 2005). These issues may be addressed using higher resolution fMRI acquisitions, measures of hemodynamic responsiveness, and methods to measure and correct signal inhomogeneities. A limitation for all neuroimaging methods involving participants with brain lesions is suboptimal spatial normalization. This issue is evident in our results as periventricular artifact where patient ventricles were larger than or displaced relative to the control group. Further advances in nonlinear spatial normalization will result in improved FAMs.

Further development of the FAM approach will also be required to address other current limitations. First, a small percentage (3%) of patient FAM models failed to converge or produced zero support vectors, indicating that the machine learning algorithm was not able to find a solution that delineated the patient from control subjects. In one case this affected a patient’s single resting state run, but in the other three cases, this affected one of multiple scan times for a given patient, suggesting that the failure to converge related to characteristics of the specific resting state run rather than characteristics of the patient lesion or brain. Another limitation is that the entirety of the densely necrotic lesion core was not detected in every person at every scan time point, perhaps due to a particularly noisy resting state run. Both of these limitations indicate room for improvement to the central FAM method.

Several avenues are available for improvement of the FAM method. Future work is needed to optimize hyperparameter selection and preprocessing procedures to achieve greater robustness to variability in the preprocessed data and more sensitivity to densely necrotic lesions. In addition, other approaches to quantifying functional aberrations in the resting signal should be considered. Here we focus on the average deviation in a patient versus a control group at a point in space, but other distributional properties of abnormal signal over the time-course of a single resting state run, such as whether abnormalities are static or fluctuate over seconds or minutes, may provide information that speaks to the nature of the anomalous signal. Speculatively, different measures (e.g., static vs. time-varying anomalies) may be sensitive to different types of dysfunction or compensation, such that multiple FAMs composed of different measures of functional anomaly may more fully capture the differences in brain function between patients and controls.

A number of potential clinical and scientific applications of FAMs should also be explored. Although we have used FAMs here to make inferences about behavior measured at a single time point, future work could associate changes in anomalous signal over time with changes in behavior over time. This may help to clarify the nature of the signal, and may also provide critical evidence regarding the brain basis of stroke recovery, the basis of day-to-day fluctuations in behavioral performance, and also the basis of behavioral response to neurorehabilitation, medications, and neuromodulation. Finally, efforts have been made to use resting BOLD data to identify functional changes in several clinical populations, often examining changes in resting state connectivity (He et al., 2007; Liu et al., 2018; Nair et al., 2015; Park et al., 2018; Ranasinghe et al., 2017); likewise, the technique described here may be applied to other populations without gross anatomical damage to identify functional anomalies in individuals with neurodegenerative disorders, psychiatric disorders, developmental disorders, or other clinical populations. Beyond clinical applications, this technique could also potentially be used to measure development or skill acquisition (e.g., literacy) or examine any source of individual difference in brain function, such as sensory experience (e.g., blind or deaf populations), multilingualism, or socioeconomic status.

While the lesion method remains a uniquely strong source of causal inference for the brain-behavior relationship, it has always been limited by its focus on anatomical damage alone. Consequently, its relevance has diminished with the explosion of functional brain imaging techniques that can address contemporary questions related to ongoing neural processes. The work we presented here shows that the lesion method can also address such questions. This represents a significant advance in a method that is thousands of years old and ensures that the study of lesions will continue to play an essential role in the neuroscientific endeavor.

## Methods

### Participants

We derive functional anomaly maps (FAMs) for a cohort of 50 left-hemisphere stroke survivors (40 ischemic, 6 hemorrhagic, 4 unknown) at least 6-months post-stroke (chronicity 49.5 ± 40.2 months (range 6.2-151.2 months); age 59.7 ± 9.3 years (range 43-78 years); 31 male, 19 female; 44 right-handed, 4 left-handed, 2 ambidextrous; education 16.1 ± 3.1 years (range 12-24 years)). Together, their strokes encompassed the middle cerebral artery distribution, with some extending into the anterior and posterior cerebral artery territories (**Figure 1**). FAMs are normed with reference to a control group consisting of 57 healthy adults (51.2 ± 21.5 years (range 19-84 years); 27 male, 28 female; 50 right-handed, 4 left-handed, 1 ambidextrous; education 16 ± 2.6 years (range 12-21 years)). Aside from the stroke events, participants had no history of psychiatric or other neurological condition. Data was collected as part of previous studies conducted in the Cognitive Recovery Lab. All subjects provided informed consent in accordance with the Georgetown University IRB. **Supplemental *Supplemental Table 1*** shows the patient demographics.

## Imaging data

### Image acquisition

Images were acquired on a 3T Siemens Magnetom Trio scanner using a 12-channel head coil at Georgetown University. T2*-weighted BOLD echo planar images were collected with the participant at rest with the following parameters: 168 + 2 initial volumes discarded; 47 axial slices in descending order; slice thickness = 3.2 mm with no gap; field of view = 240 × 240 mm; matrix 64 × 64; repetition time (TR) = 2,500 ms; echo time (TE) = 30 ms; flip angle = 90°; voxel size = 3.2 × 3.2 × 3.2 mm. For a subset of participants, runs were 239 volumes long, which was cut the 168 to make them comparable to other runs. For anatomical reference and lesion tracing, T1-weighted MPRAGE structural images (voxel size = 0.98 × 0.98 × 1.0 mm) were acquired.

### Multiple sessions

We included in our analyses multiple resting state runs acquired at different times for participants as available. In total, 132 resting state runs across 50 subjects were analyzed. Of the 36 subjects with multiple resting state scans, five had two sessions, 25 had three sessions, four had four sessions, one had six sessions, and one had eight sessions. The time between first and last scan sessions was 5.2 ± 9.8 months (range 0.4-63 months; median = 3.3 months). For patients with multiple sessions, the first structural scan served as a shared anatomical reference.

### Anatomical lesion tracings

Patient lesions were traced based on T1-weighted anatomical images by a trained neurologist (P.E.T.) using ITK-snap (Yushkevich et al., 2006), and were used as cost-function masks during normalization of patient anatomical scans (Brett et al., 2001).

### Image preprocessing

Imaging data were preprocessed using the CONN toolbox version 18a (Whitfield-Gabrieli and Nieto-Castanon, 2012) and SPM12 (Friston, 2003) using the default pipeline for volume-based analyses. Specific preprocessing steps included functional realignment and unwarping, image centering, and slice-timing correction. Scans with global signal *Z* > 9 or subject-motion of >2mm were rejected as outliers. Data were smoothed by spatial convolution with an 8mm FWHM Gaussian kernel. A nuisance model was used to remove confounders, including the 6 realignment parameters estimated during correction for minor head movement and their first-order derivatives, the effect of rest and its first-order derivative, and scrubbing parameters for outlier volumes. The residuals were linearly detrended and band-pass filtered (0.008-0.09 Hz). Cerebrospinal fluid signal and white matter signal were not included as covariates of the nuisance model to ensure that we did not inadvertently remove signal that distinguished abnormal regions, such as areas of encephalomalacia filled with cerebrospinal fluid. Structural scans for each subject were segmented and normalized to the Montreal Neurological Institute (MNI) average of 152 brains in SPM12. During normalization, patient brains were cost-function masked with manually-traced anatomical lesion masks. Functional time-series and anatomical lesion tracings for each participant were then normalized using these warp fields.

### Machine learning approach

In this study we use support vector regression (SVR), a machine-learning approach to multiple regression (Drucker et al., 1996) adapted from the support vector machine (Cortes and Vapnik, 1995). Specifically, our analyses utilize epsilon-insensitive SVR (ε-SVR) implemented in MATLAB 2018a. As its name suggests, SVR behaves comparably to regression but importantly it uses a machine learning algorithm for parameter estimation, which allows it to be robust to many collinear predictors (here, individual voxels). From an algorithmic standpoint, the technique is also innately similar to the process of manual lesion segmentation, in which an experimenter classifies voxels by tracing out a boundary line between damaged and spared tissue. Similarly, the classic SVM finds an optimal decision boundary that separates classes of labeled data. In practice, the input data features are transformed into a high-dimensional space in which an optimal hyperplane is estimated. The resulting model is comprised of a subset of training points, referred to as support vectors, which constrain this estimated hyperplane. Although a trained model is usually then used to classify novel, unobserved data sets, here we back-project the model into the original dataspace to create a pseudodataset in which we can visualize voxels that play an important role in defining the hyperplane solution.

We use SVR as opposed to SVM for functional anomaly mapping because it treats differences between patients and controls as falling on a continuum rather than unnecessary assuming they are mutually exclusive classes that are categorically distinct. SVR is a mathematical reformulation of the SVM classification problem in which the hyperplane can be thought of as falling through the center of the solution like a line of best fit, rather than separating classes.

### Functional anomaly map (FAM) derivation

We used a novel procedure to derive a FAM for each resting state run of each patient. This procedure involved comparing a normalized, pre-processed 4D resting state time-series from a single patient to a control group with the following procedure: All data were combined into a single feature matrix with columns corresponding to subjects and rows corresponding to points in space and time. These data comprised the training features of an SVR model with a contrast matrix specified as patient status coded with an indicator variable. SVR-β values from the trained model were back-projected into 4D (see equation 8 of Zhang *et al.*, 2014). Each back-projected 3D volume was then *Z*-scored to eliminate uninterpretable volume-to-volume variability. Finally, a single 3D image was computed as the average absolute *Z* value down the fourth dimension.

The absolute value was used because the sign of the *Z*-scores have no straightforward interpretation, owing to the fact that the sign of the original resting state data naturally fluctuates.

This procedure yielded a single 3D statistical map of functional anomaly that can be overlaid onto a brain in standard space. The value at each voxel was an average *Z*-score, reflecting the accumulation of statistical deviations from the control group discovered by the SVR in the individual patient’s resting state run.

Functional anomaly mapping analyses employed a linear kernel with the following hyperparameters: epsilon = 0.1; box constraint = 1; kernel scale = auto; standardize = true; outlier fraction = 5%; algorithm = ISDA. Kernel scale was chosen by a heuristic subsampling procedure implemented in MATLAB. Our analyses were conducted with 4mm^3^ voxels within a brain mask (Holmes et al., 1998).

### Validation of FAMs through localization of scores with ground-truth localization

For analyses validating the FAM signal, scores with ground-truth localizations were generated based on the anatomical lesion load for each patient within 132 left-hemisphere atlas parcels (Shen et al., 2013). For each score, an SVR-LSM analysis (see Lesion-Behavior Mapping below) was performed using FAMs as lesion maps. Localization accuracy was calculated as the Euclidean distance between the center-of-mass of the source atlas parcel and the center-of-mass of its SVR β-map solution (thresholded at 7). To determine whether localization accuracy was greater than chance, we compared the distribution of actual solution distances to the distribution obtained if solution distance was randomly related to seed. This null distribution consisted of the distances between each seed’s center-of-mass and the center-of-mass for all solutions. To assess whether right-hemisphere FAM signals could detect functional lesions in the homotopic region of the left hemisphere, we generated another set of 132 scores using the average FAM for each patient in each of the 132 left-hemisphere atlas parcels, and performed SVR-LSM analyses for each score using only the right hemisphere of the FAMs as lesion maps. For each score, the localization accuracy was calculated as above, ignoring the sign of the *x* coordinate.

### Behavioral scores for lesion-behavior mapping

Four behaviors with known patterns of localization were examined for lesion-behavior mapping using FAMs and anatomical lesion maps. Auditory comprehension impairment was measured using the Yes/No Questions subtest on the Western Aphasia Battery (WAB) (Shewan and Kertesz, 1980). Phonological processing deficits were measured using an in-house pseudoword repetition test. Speech fluency deficits were measured by the mean length of utterance (MLU) during the Picnic Scene picture description from the Western Aphasia Battery and/or the Cookie Theft Picture from the Boston Diagnostic Aphasia Examination (Goodglass et al., 2001). To control for the contribution of lexical-retrieval deficits during the picture description task, we included as a covariate the accuracy on a spoken picture naming test (Philadelphia Naming Test) (Roach et al., 1996). Upper-limb motor impairment was measured using the right-hand pinch-strength score on the Motricity Index (Demeurisse et al., 1980).

### Lesion-behavior mapping

Anatomical lesion-behavior mapping was conducted with 2.5 mm^3^ voxels using SVR-LSM implemented in our MATLAB toolbox using default hyperparameters (DeMarco and Turkeltaub, 2018; Zhang et al., 2014). Lesion volume was regressed out of both the behavior and voxelwise data as a covariate of no interest using a nuisance model. Functional lesion-behavior mapping was conducted with 4 mm^3^ voxels invoking modified versions of these same toolbox routines. Functional lesion-behavior mapping analyses employed the following hyperparameters: epsilon = 0.1; box constraint = 1; kernel scale = auto; standardize = true; outlier fraction = 5%; algorithm = ISDA. All results were thresholded voxelwise at *P* < .005, and corrected for multiple comparisons based on cluster extent at *P* < .05, based on 10,000 permutations.

## Acknowledgements

The authors would like to thank Elizabeth Lacey, Mackenzie Fama, Zainab Anbari, and Kate Spiegel for data collection. The research was supported by the Doris Duke Charitable Foundation (grant #21012062), NIH/NCATS via GHUCCTS (KL2TR000102 and TL1TR001431), and by the NIDCD (R01DC014960).

## Authors Contributions

Conceptualization, A.T.D. and P.E.T.; Methodology, A.T.D. and P.E.T.; Investigation, A.T.D. and P.E.T.; Formal Analysis, A.T.D.; Writing – Original Draft, A.T.D.; Writing – Review & Editing, A.T.D. and P.E.T.; Funding Acquisition, P.E.T. and A.T.D.; Supervision, A.T.D.

---

**Supplemental Figure 1.**
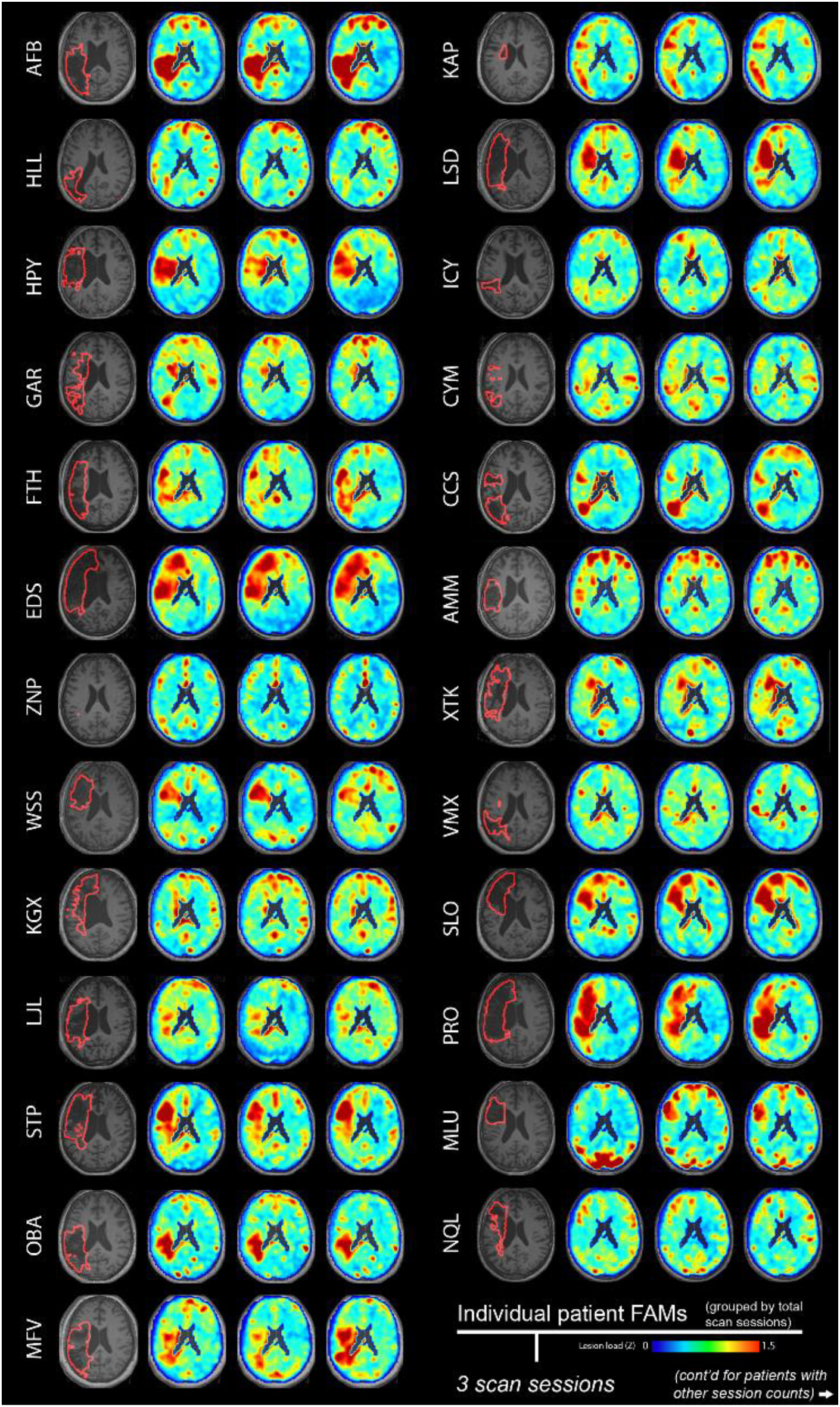
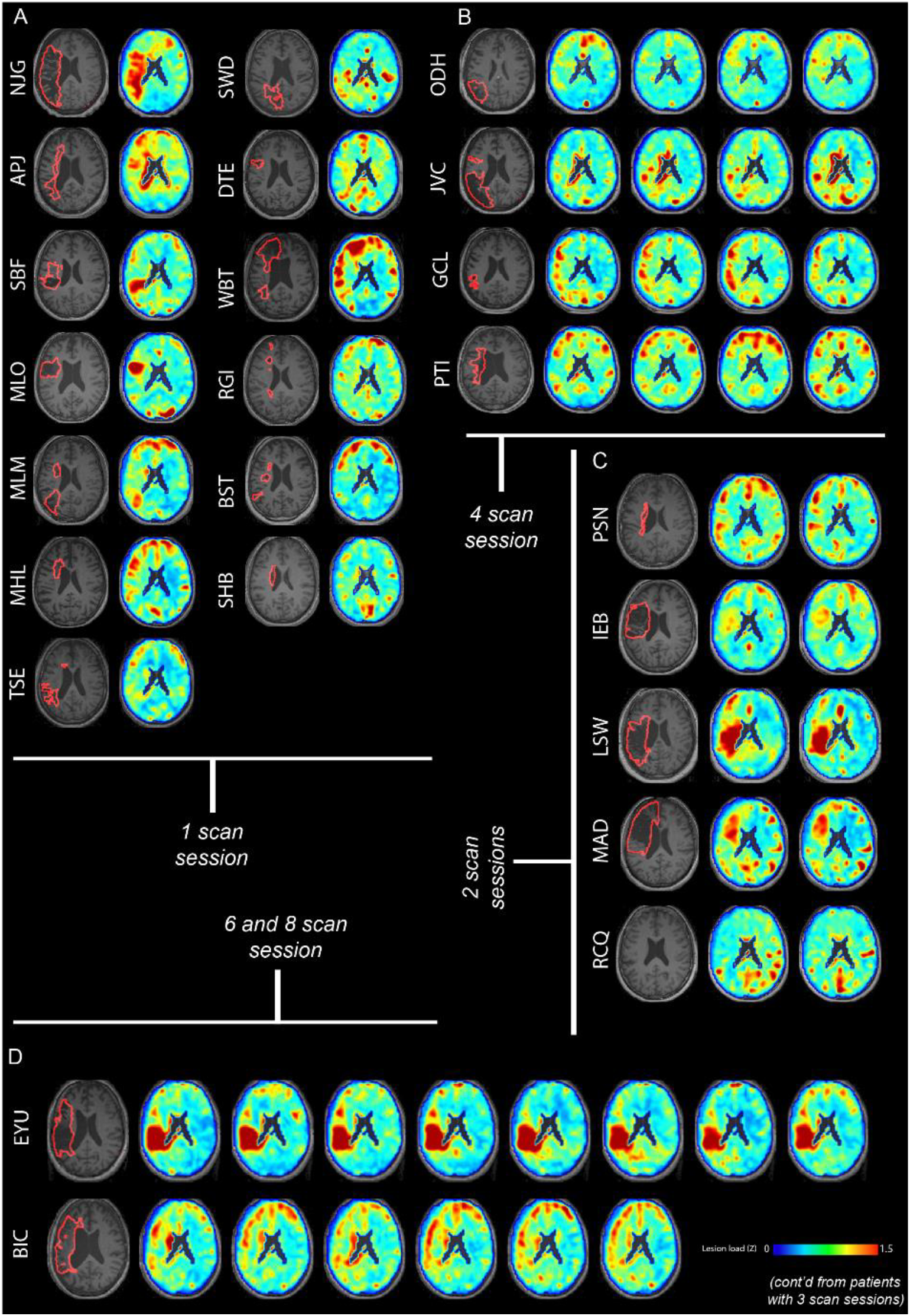
FAMs for each patient at all available time points. Part 1 shows FAMs for each patient with three scan sessions. Part 2 (A-D) shows FAMs for each patient with two, four, six, or eight scan sessions. For each patient, the left-most image shows the anatomical scan with the anatomical lesion outlined in red. All FAMs are shown in a jet color scale.

**Supplemental Figure 2.**
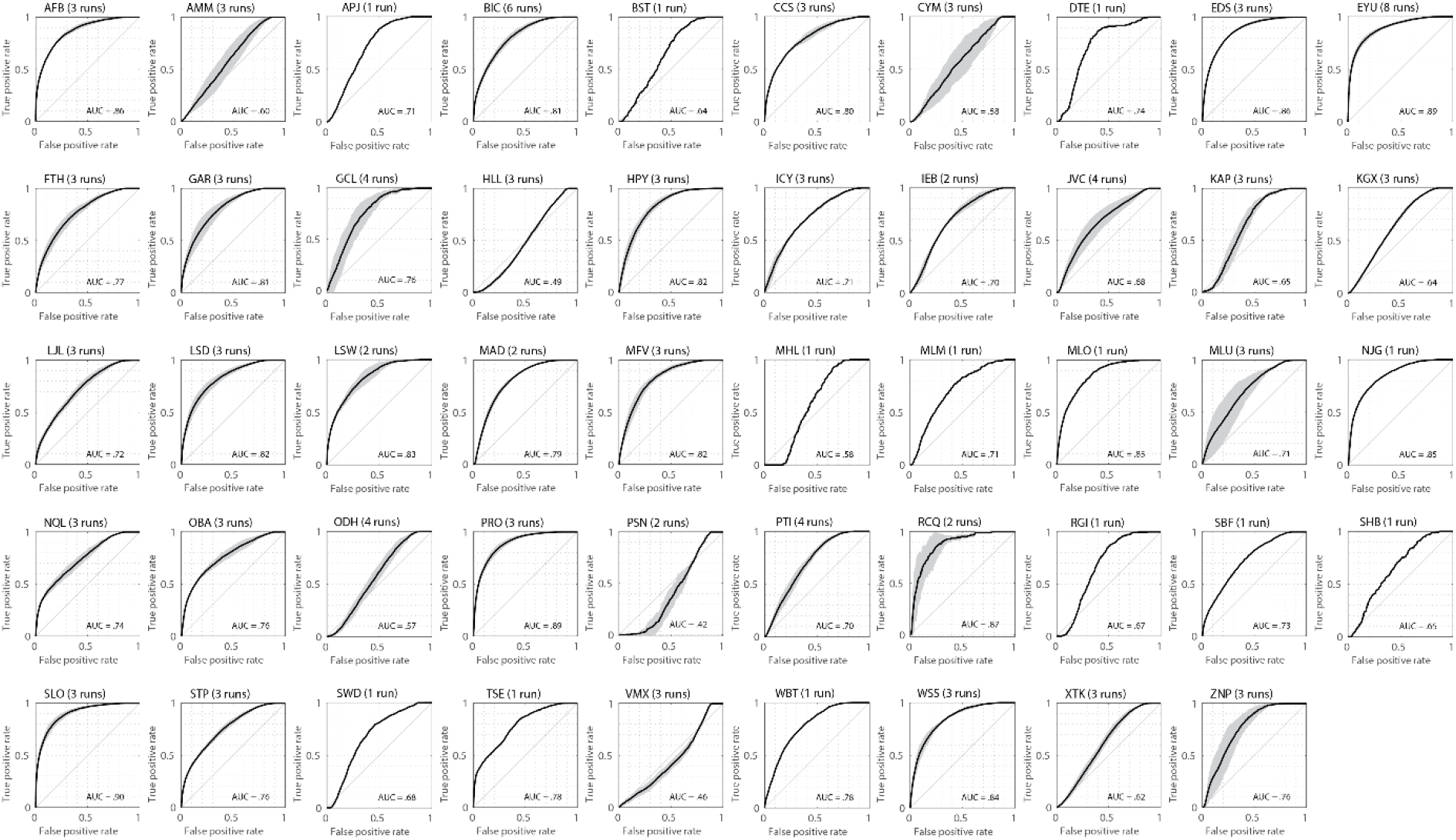
For each patient, ROC curve and average AUCs for the accurate classification by FAMs of manually-traced tissue as damaged or spared. Each plot shows the participant code, the number of runs that contribute to the plot for that subject, the average ROC curve (black), and the standard deviation around the curve (dark gray) for participants with multiple scan sessions.

**Supplemental Table 1.**
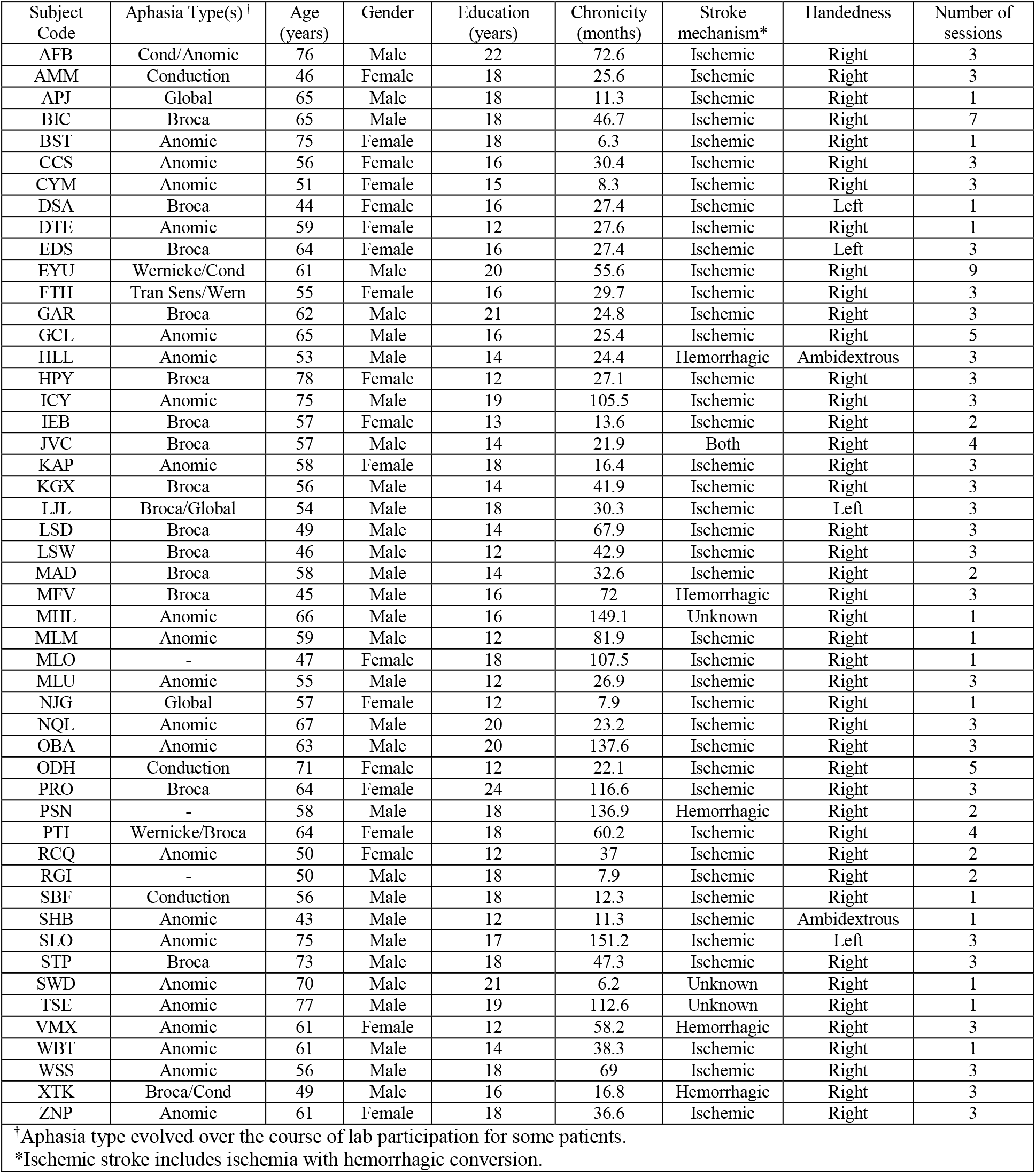
Patient cohort demographics in this study.

